# Strain-Specific Human Natural Killer Cell Recognition of Influenza A Virus

**DOI:** 10.1101/148528

**Authors:** Lisa M. Kronstad, Christof Seiler, Rosemary Vergara, Susan P. Holmes, Catherine A. Blish

## Abstract

**Abstract:** Innate Natural killer (NK) cells employ an array of surface receptors to detect ‘altered self’ induced by infection or malignancy. Despite their decisive role in early antiviral immunity, the cellular mechanisms governing if or how they discriminate between viral infections remain unresolved. Here, we demonstrate that while human NK cells are capable of reducing infection levels of distinct influenza A strains, the A/California/07/2009 (pH1N1) strain induces a significantly more robust IFN-γ response than A/Victoria/361/2011 (H3N2) and all other strains tested. This surprising degree of strain specificity results in part from the inability of the pH1N1 strain to downregulate the activating ligands CD112 (Nectin-2) and CD54 (ICAM-1) as efficiently as the H3N2 strain, leading to enhanced NK cell detection and IFN-γ secretion. A network analysis of differentially expressed transcripts identifies the interferon α/β receptor (IFNAR) pathway as an additional, critical determinant of this strain-specific response. Strain-specific downregulation of NK cell activating ligands and modulation of type I IFN production represents a previously unrecognized influenza immunoevasion tactic and could present new opportunities to modulate the quality and quantity of the innate antiviral response for therapeutic benefit.

**One Sentence Summary:** Human natural killer cells distinguish between Influenza A strains using a combinatorial cytokine priming and receptor-ligand signaling mechanism.

## Introduction

NK cells are innate lymphocytes that defend from viral infection by killing infected cells and regulating the subsequent adaptive immune response (*1*, *2*). While B and T cells use somatically recombined antigen-specific receptors, NK cells rely on a complex array of germline-encoded inhibitory and activating receptors to regulate their effector functions (*3*-*5*). NK cell inhibitory receptors recognize HLA class I molecules, holding NK cell activity in check upon encounter with healthy cells (*6*, *7*). If infected or malignant cells downregulate HLA class I expression, these cells fail to deliver an inhibitory signal through NK cell inhibitory receptors, which include killer immunoglobulin (Ig)-like (KIRs), CD94-NKG2A heterodimers, and ILT2 receptors (*8*, *9*). NK cell activating receptors recognize diverse cellular ligands upregulated by infection or malignancy, flagging these abnormal cells as potential NK cell targets by overruling inhibitory signals (*10*). Major NK cell activating receptors are generally non-HLA restricted and include the Fc receptor CD16, NKG2D, NKG2C, NKp30, NKp44, NKp46, NKp80, the co-stimulatory receptors CD226 (DNAX-accessory molecule-1, DNAM-1), CD244 (2B4), and a variety of adhesion molecules including LFA-1 (CD11a/CD18 heterodimer) and CD2 (*4*, *10*). In addition to these direct receptor-ligand interactions, NK cells are responsive to soluble activators such as cytokines through expression of IL-12, IL-15, IL-18, IFN-α and -γ receptors (*11*, *12*). Collectively, appropriate integration of receptor-ligand and cytokine-mediated signals is integral for NK cells to spare normal cells while maintaining their capacity to rapidly and robustly respond to abnormal target cells (*1*, *4*). Yet how these distinct receptors and cytokines work in a combinatorial fashion to recognize specific viruses remains unknown.

Influenza infection is a relevant system to dissect the mechanisms by which NK cells recognize and respond to distinct viral strains, as the virus undergoes constant evolution, giving rise to seasonal epidemics and periodic pandemics. Evidence from both murine and human studies indicates a role for NK cells in regulating the outcome of influenza infection *in vivo*. In mice, NK cells represent a substantial proportion of immune cells in healthy lungs, and peripheral NK cells traffic to the lung following influenza infection *(13-16)*. Mice depleted of NK cells by treatment with anti-asialo GM1 or anti-NK1.1 antibodies had increased morbidity and mortality and failure to induce influenza A virus-specific cytotoxic T lymphocytes after sublethal influenza challenge *(17, 18)*. A specific role for the NK cell cytotoxicity receptor NCR1, a homolog of the human NKp46 natural cytotoxicity receptor, has been suggested based on its binding to influenza hemagglutinin *(19, 20)*. One strain of NCR1-deficient mice was highly susceptible to influenza infection *(21, 22)*, while another strain, which lacked surface expression of NCR1, displayed both a hyper-responsive NK cell phenotype and increased resistance to influenza infection. Thus, the precise contributions of NK cell NCR1 expression on influenza outcomes in mice remain unclear. Separate studies report that NK cells may play a detrimental role during influenza infection, as IL-15^-^/^-^ mice, which lack NK cells, and mice depleted of NK1.1-expressing cells had less pulmonary inflammation and reduced mortality after lethal-dose influenza infection *(23, 24)*. Collectively, these data suggest a dual role for NK cells during influenza infection, either contributing to immunopathology or conferring protection, depending on the viral dose and murine model. This dual potential, to enhance or hinder recovery from infection, emphasizes the need to understand the elements governing the quality and quantity of the NK cell response to influenza.

In humans, several studies suggest NK cells may contribute to the outcome of infection. As with the murine infection model, human NK cells represent a substantial portion of immune cells in healthy lungs and are further recruited to the lung during influenza infection *(25)*. During the 2009 H1N1 pandemic, 100% of subjects with severe infection developed NK cell lymphopenia, compared to 13% of mild cases *(26, 27)*. Earlier reports found that patients with severe influenza infection had a near complete lack of pulmonary NK cells *(28, 29)*. Thus, NK cell deficiency is associated with severe influenza infection in humans.

It is not clear which specific receptors contribute to influenza recognition by human NK cells. Influenza vaccination is associated with an expansion in 2B4^+^ NK cells, a decline in the frequency of NKp46^+^ NK cells, and enhancement of IL-2-dependent NK cell responses to exogenous cytokines *(30-32)*. Influenza hemagglutinin (HA) directly interacts with NKp46 in a sialic acid-dependent manner *(33, 34)*, although the consequence of NKp46-HA interactions during *in vivo* infection is unknown. *In vitro*, NKp46 and NKG2D contribute to the response of human NK cells to dendritic cells infected with the influenza A/Panama/2007/1999 strain (*35*). In another study, the NK cell IFN-γ response to *in vitro* influenza A virus stimulation was dependent upon IL-2 produced by T cells (*36*). We previously observed that human NK cells mount distinct IFN-γ immune responses to purified A/California/07/2009 (pH1N1) and A/Victoria/361/2011 (H3N2) strains in the presence of T cells *in vitro (37)*, calling into question the mechanisms governing human NK cell recognition of antigenically divergent influenza strains.

Here, we report that NK cells distinguish viral strains, mounting a far more robust IFN-γ response to the 2009 pandemic H1N1 strain than to the 2011 H3N2 and several other strains. To identify the source of differential NK cell activation, we used a 38-parameter mass cytometry approach to profile influenza A-mediated modulation of NK cell ligand expression at single-cell resolution and RNA-sequencing to identify a unique transcriptional signature of pH1N1-vs. H3N2-infected cells *(38)*. Specifically, we provide evidence that modulation of NK cell activating ligands and the interferon α/β receptor (IFNAR) pathway governs human strain-specific innate responses to influenza A infection.

## Results

### NK cells recognize antigenically divergent influenza A viral strains

We established a co-culture system to assess the ability of human NK cells to recognize and respond to influenza A infection in autologous monocytes (**Fig. S1A**) (*36*). To investigate the strain specificity of the NK cell response, NK cells were co-cultured with monocytes either mock-treated or infected at an MOI of 3 with A/Victoria/361/2011, A/California/07/2009 (pH1N1), A/Bayern/7/1995, A/Beijing/32/1992, A/Hong Kong/1/68 or A/Melbourne/1/1946. Each strain infects monocytes and drives a NK cell functional response, as measured by expression IFN-γ and CD107a, a marker of degranulation (**Fig. 1A-C**). The magnitude and quality of NK cell response does not map to subtype (**Fig. 1A-C**). A significantly greater proportion of NK cells secrete IFN-γ in response to the pH1N1 strain than to all other strains tested (**Fig. 1A**).

**Fig. 1.**
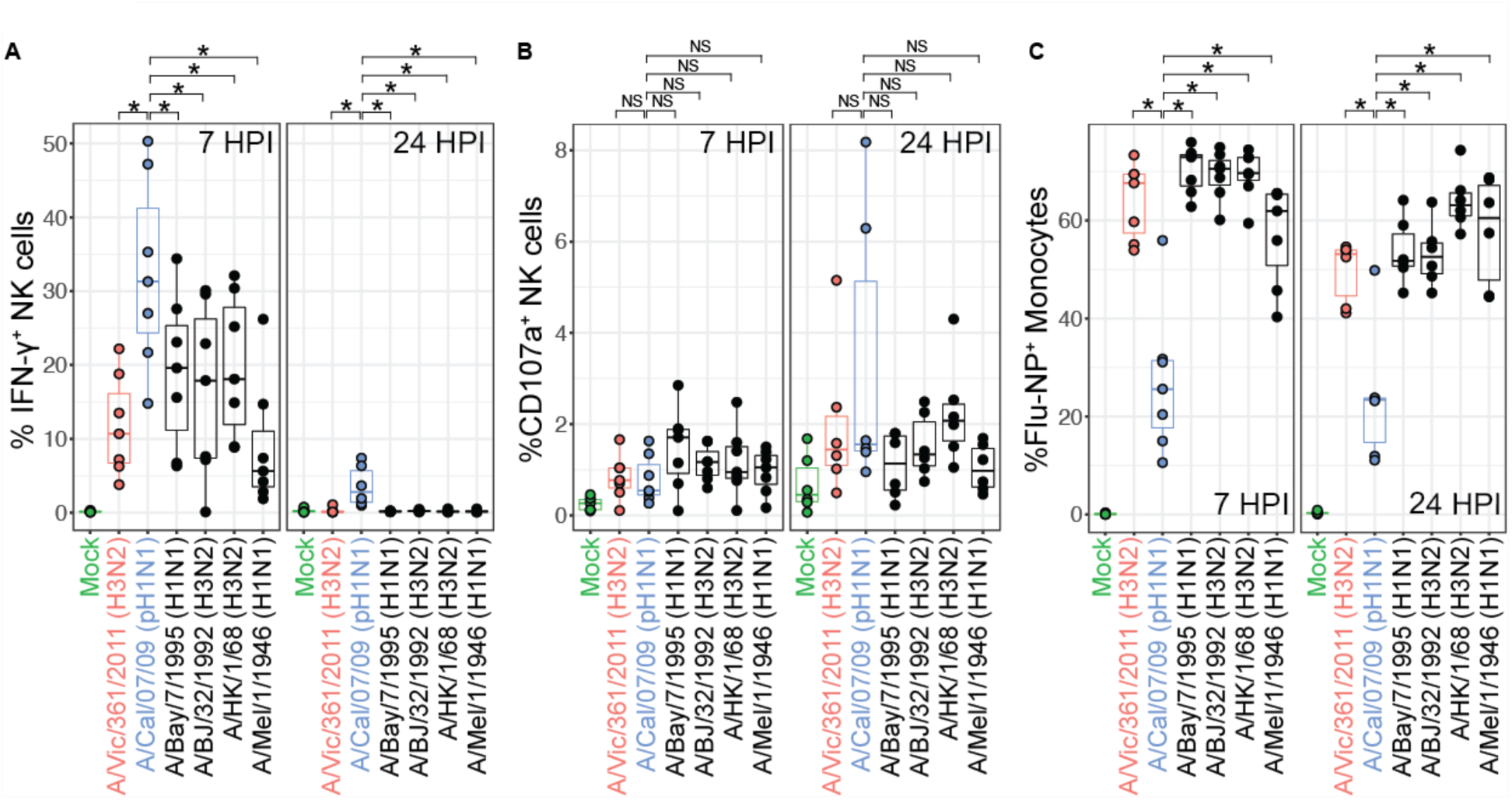
NK cells secrete IFN-γ and degranulate in response to autologous monocytes infected with six influenza A strains. (**A**) NK cell IFN-γ production, (**B**) CD107a expression and (**C**) monocyte infection levels (Flu-NP) to the indicated influenza A strains at 7 HPI (*n*=7) or 24 HPI (*n*=6). NS *P* > 0.05, \**P* < 0.05, Wilcoxon signed-rank test.

### pH1N1 infection triggers enhanced NK cell IFN-γ production compared to H3N2

We next selected A/Victoria/361/2011 (H3N2) and A/California/07/2009 (pH1N1), the most recently circulating strains, to elucidate the source of the strain-specific NK cell IFN-γ response. The H3N2 strain infects a significantly greater frequency of monocytes than the pH1N1 strain at an identical multiplicity of infection (MOI), though neither strain infects a significant proportion of NK cells (**Fig. S1B-C**). Monocytes infected with both pH1N1 and H3N2 induce a significant frequency of NK cells to secrete IFN-γ at 7 hours post-infection (HPI) (**Fig. 1A, 2A**). The pH1N1 strain elicits an ~7-fold greater frequency of NK cells to secrete IFN-γ than does the H3N2 strain at 7 HPI (pH1N1 median=21.6%, H3N2 median=3.21%, *P*=0.0001) (**Fig. 2A**). This difference holds true at 24 HPI, although responses are lower in response to both strains (pH1N1 median=1.58%, H3N2 median=0.15%, *P*=0.004) (**Fig. 2A**). Infected monocytes are required for activation, as purified NK cells exposed to virions do not secrete IFN-γ (**Fig. 2A**). NK cell activation is independent of T cell help, as robust NK cell activation was observed following stringent T cell depletion and was not enhanced upon re-addition of T cells (**Fig. 2A and S2**). Moreover, the increased response to the pH1N1 strain is observed regardless of the infection level observed in monocytes, as it persists even after increasing the infection levels in pH1N1 to reach equivalence with H3N2 infection levels (**Fig. S3**). IFN-γ is also secreted into the supernatant and transcribed at significantly greater levels in response to pH1N1 compared to H3N2-infected monocytes (**Fig. 2B-C**).

**Fig. 2.**
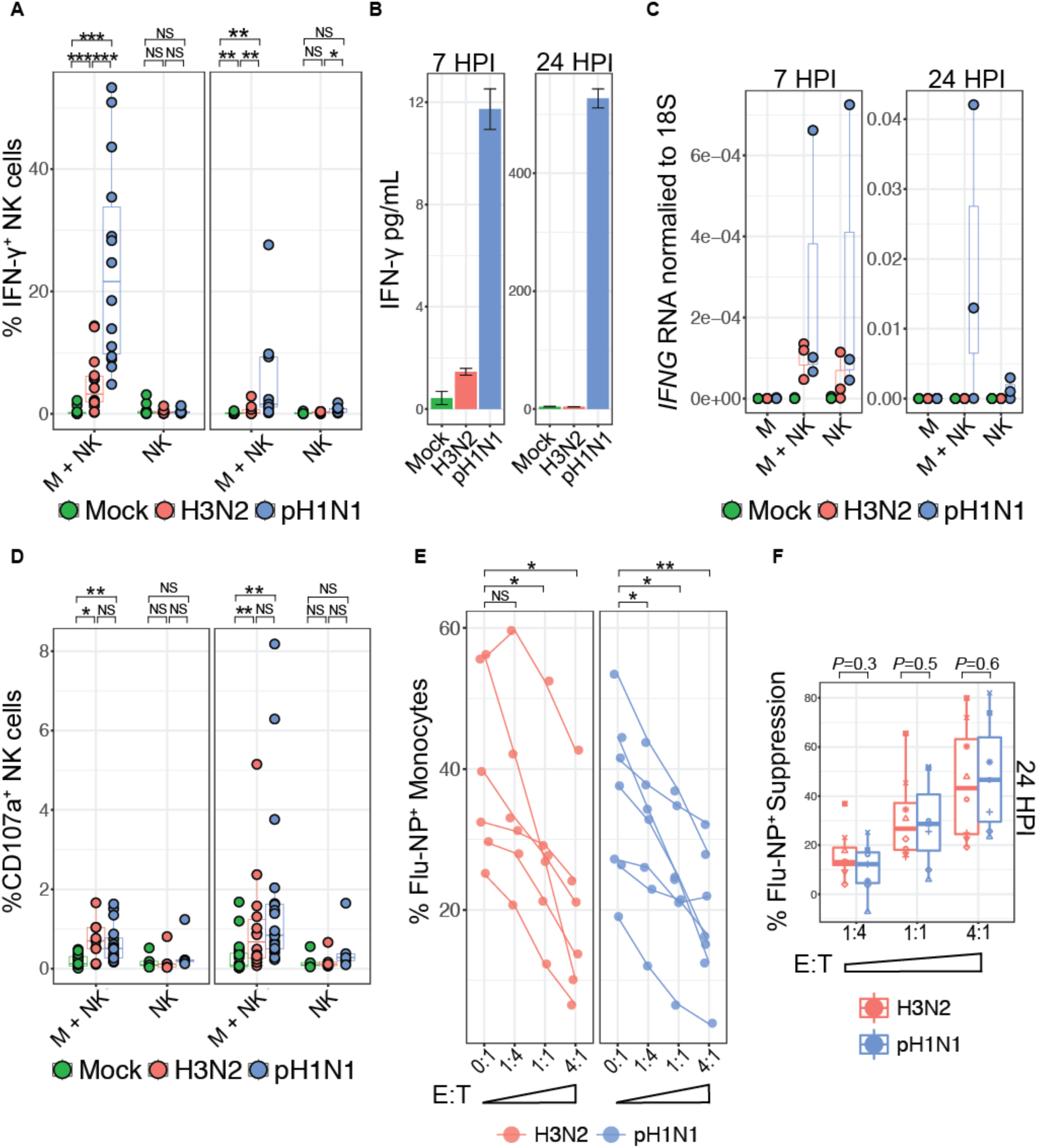
NK cells suppress infection and secrete IFN-γ in response to autologous pH1N1 and H3N2 infected monocytes. **(A)** Percent IFN-γ^+^ NK cells post exposure to mock, H3N2- or pH1N1-infected monocyte or virions at 7 or 24 HPI assessed by intracellular cytokine staining. **(B)** Supernatants harvested from mock, H3N2- or pH1N1 monocyte – NK cell co-cultures (without brefeldin/monesin) after 7 or 24 hours were subjected to 63-plex Luminex^®^ assay. Average of technical duplicates from a representative donor with standard deviation is plotted. **(C)** *IFNG* transcript levels normalized to 18S in pure monocyte (M), monocyte – NK cell co-culture (M+NK) or pure NK cells (NK) after either mock treatment, H3N2 or pH1N1 exposure (MOI=3). Values represent average of technical triplicates (*n*=3). (**D**) Percent CD107a^+^ NK cells post exposure to mock, H3N2- or pH1N1-infected monocyte or virions at 7 or 24 HPI with an anti-CD107a antibody added for the last four hours of co-culture and assessed by flow cytometry. (**E**) The percent Flu-NP^+^ monocytes measured 24 HPI with H3N2 (*n=6)* or pH1N1 (*n*=7) (MOI=3) and cultured at an effector to target (E: T) cell ratio of 0:1, 1:4, 1:1 or 4:1. (**F)** Boxplot representation of percent viral suppression at each E:T ratio (*n*=9). NS *P* > 0.05, \**P* < 0.05, ** *P* < 0.005, *** *P* < 0.0005, Wilcoxon signed-rank test.

### NK cells suppress pH1N1 and H3N2 infection levels

Co-culture of NK cells with infected monocytes leads to expression of CD107a and significantly reduces the frequency of infected monocytes (**Fig. 2D and S4A**). Significant individual variation is observed in the magnitude of initial infection levels and in the reduction of infection (**Fig. 2E and S4A**). Direct comparison of the NK cell viral suppression across individuals indicates that NK cells achieve ~40% suppression of both strains (**Fig. 2E-F**). When infection was allowed to proceed for 7 hours, NK cell reduced infection levels to a lesser extent (**Fig. S4A-B**). Transcription of the influenza matrix RNA gene segments is also suppressed upon the addition of NK cells, reaching nearly 100% suppression of viral transcription for both strains (**Fig. S4C-D**). Collectively, these data indicate that NK cells can suppress viral protein and mRNA levels in cells infected with pH1N1 and H3N2 influenza strains.

### Direct contact is required for NK cells to sense infection and suppress infection levels

To identify whether receptor-ligand interactions are required for NK cell responses to influenza infection, NK cells and monocytes were co-cultured in a *trans*-well system (**Fig. 3A**). Elimination of direct NK cell-monocyte contact completely abrogates the NK cell IFN-γ response to both H3N2 and pH1N1-infected monocytes (**Fig. 3B**). Secreted IFN-γ protein levels are also dramatically decreased upon elimination of cell-cell contact (**Fig. 3C**). Elimination of direct contact diminishes the ability of NK cells to express CD107a, a marker of cytolysis, and to suppress viral infection levels (**Fig. 3D-E**). Collectively, the impairment of NK cell effector functions upon physical separation is consistent with receptor-ligand interactions being necessary for the NK cell anti-influenza response.

**Fig. 3.**
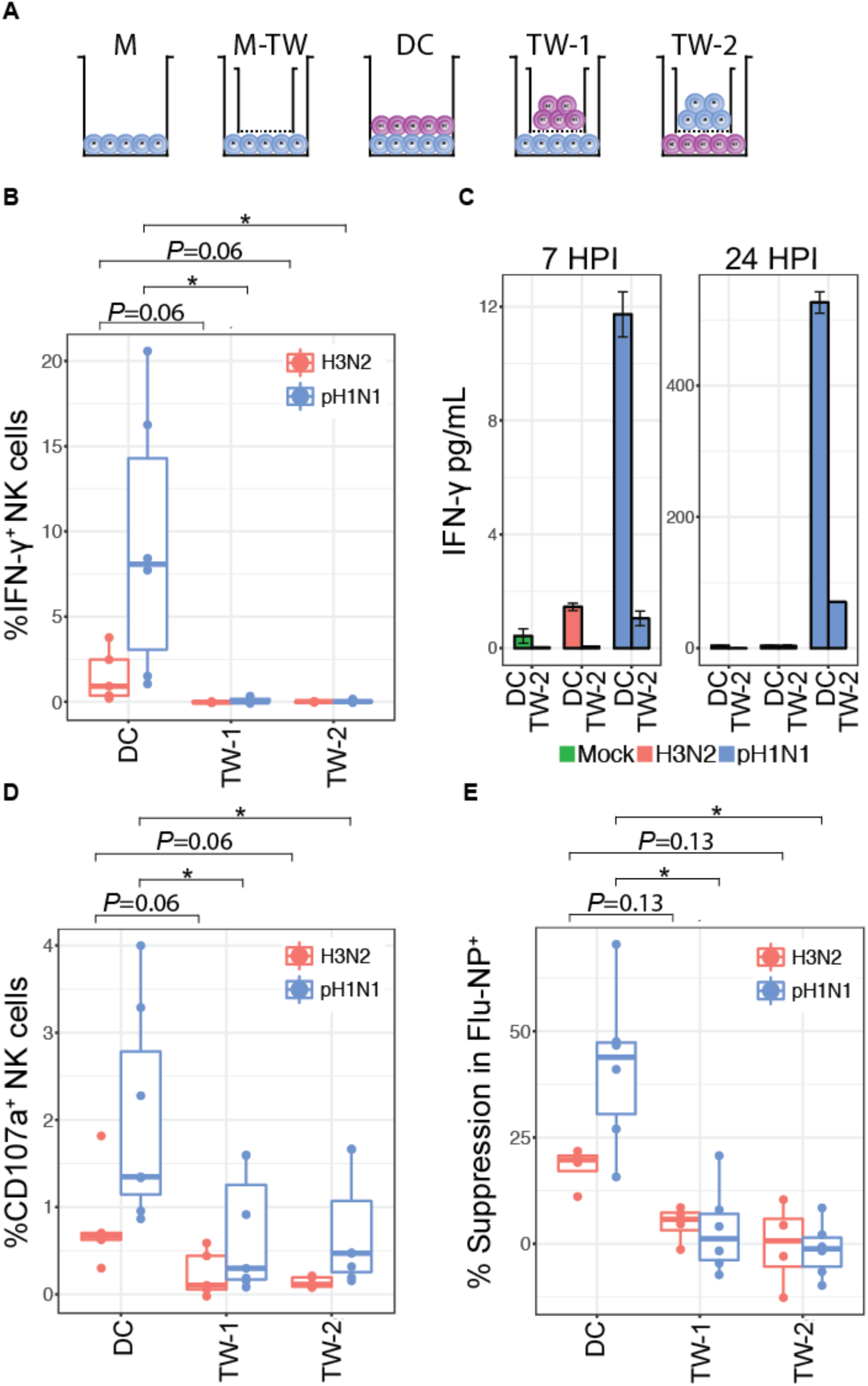
NK cells require direct contact for viral suppression, CD107a expression, and IFN-γ production. **(A)** Monocytes were cultured either alone (M), with a *trans-*well (M-TW), in direct contact with NK cells (DC), with monocytes seeded in dish and NK cells seeded in the *trans*-well (TW-1), or monocytes seeded in the *trans*-well and NK cells seeded in the dish (TW-2). **(B)** Percentage of IFN-γ^+^ NK cells after 6 hr co-culture with infected monocytes; H3N2 (*n*=5), pH1N1 (*n*=6). (**C**) IFN-γ concentrations in supernatants harvested from mock, H3N2-infected monocyte or pH1N1-infected monocyte co-culture with NK cells after 7 or 24 hours from DC or TW-2 condition assessed by Luminex® assay. Average of technical duplicates from a representative donor. Error bars represent standard deviation. **(D)** Percentage of CD107a^+^ NK cells after 23 hr co-culture with infected monocytes; H3N2 (*n*=5), pH1N1 (*n*=6). (**E**) Percent viral suppression was calculated based on the reduction in Flu-NP^+^ H3N2-infected monocytes (*n*=4) or pH1N1-infected monocytes (*n*=6). Mock-treated values were subtracted from infected measurements. \**P* < 0.05, Wilcoxon signed-rank test.

### Influenza infection modulates inhibitory and activating ligands on H3N2 and pH1N1-infected cells

To identify the receptor-ligand interactions required for the NK cell response to influenza-infected monocytes, we designed a 38-parameter mass cytometry panel to quantify the expression of ligands that NK cells might use to distinguish infected cells (**Table S1 and Fig. S5-7**). A generalized linear mixed model (GLMM) was used to identify the NK cell ligands modulated by influenza infection of monocytes. At 24 HPI, expression of several markers is predictive of H3N2 infection, including CD95, Flu-NP, CD11b, HLA-E, HLA-DR, HLA-C, CD33 and PAN-HLA (**Fig. 4A**). Intracellular levels of the influenza Flu-NP are strongly predictive of influenza infection, a reassuring validation of this unbiased approach. Several markers are predictive of mock-treated monocytes, including LFA-3, CD14, CD155 (PVR), IFN-α, and CD111 (**Fig. 4A**). A principle component analysis (PCA) separates mock- from H3N2-infected monocytes and identifies markers that contribute to the observed variance to differing degrees (**Fig. 4B**). Consistent with the results from our GLMM, variation in CD14 and CD155 (PVR) distinguish mock-treated samples (**Fig. 4B**). CD14 is downregulated on influenza-infected cells (**Fig. S6B**), and these data suggest that the H3N2 virus may downregulate CD155 in an attempt to escape NK cell responses. CD155 is the cognate ligand for CD226 (DNAX accessory molecule-1, DNAM-1), an NK cell activating receptor.

**Fig. 4.**
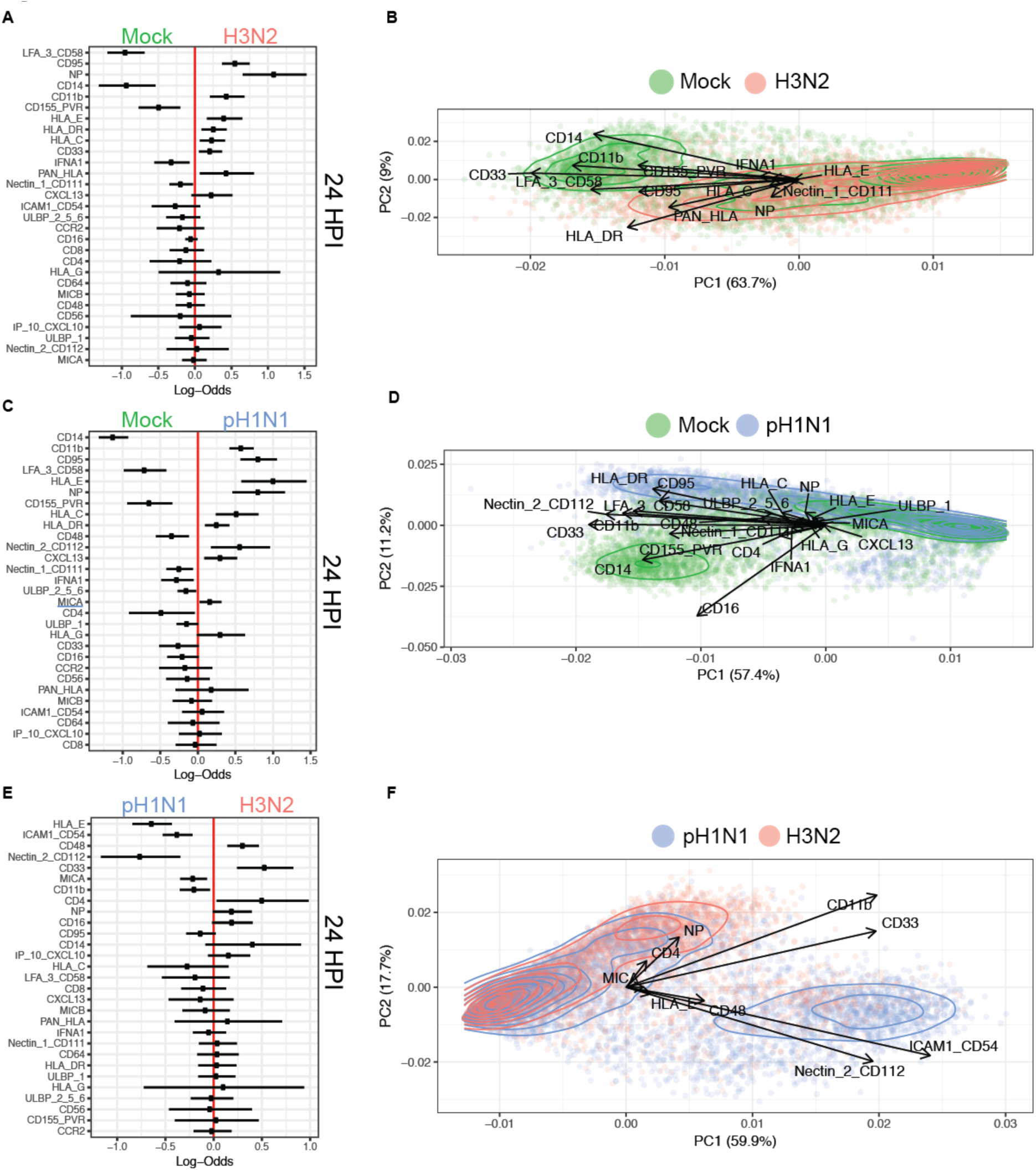
Identification of influenza-mediated modulation of NK cell inhibitory and activating ligands by mass cytometry and GLMM analysis. (**A**) Generalized linear mixed model (GLMM) to identify markers predictive of mock vs. H3N2 infection. Log-odds are logarithm of ratios of the probability that a cell is H3N2-infected over the probability that a cell is mock-infected at 24 HPI. An increase in the parameter coefficient corresponds to the strength of the classification power, with the 95% confidence interval from sampling error represented by line surrounding the point estimate. The confidence intervals are widened using Bonferroni’s method. The reported p-values are controlled using Benjamini-Hochberg’s false discovery rates. (**B**) PCA of individual cells colored by mock vs. H3N2 infection at 24 HPI, with the vectors driving variance displayed. (**C**) GLMM prediction of mock vs. pH1N1 infection at 24 HPI. (**D**) PCA of mock vs. pH1N1 infection at 24 HPI. (**E**) GLMM prediction of pH1N1 vs. H3N2 infection at 24 HPI. (**F**) PCA of pH1N1 vs. H3N2 infection at 24 HPI.

We next assessed pH1N1 modulation of NK cell ligands on infected monocytes. By 24 HPI, CD11b, CD95, HLA-E, Flu-NP, HLA-C, HLA-DR, CD112, CXCL13 and MICA are predictive of pH1N1 infection, while CD14, LFA-3, CD155, CD48, CD111, IFN-α, ULBP2/5/6 and CD4 predict mock-treated monocytes (**Fig. 4C**). PCA visualization distinguishes mock-from pH1N1-infected monocytes, though the drivers of this variance are distinct from those observed during H3N2 infection (**Fig. 4D**). While CD14 and CD155 continue to be predictive of mock-treatment, expression of several HLA molecules (HLA-DR, HLA-C, HLA-E) in addition to Flu-NP and CD95 are the most significant drivers of the variance between pH1N1 infection and mock (**Fig. 4D**). Similar findings are observed at 7 HPI, though differences are subtler with shorter infection periods (**Fig. S8-9**).

To investigate how pH1N1 elicits an enhanced NK cell INF-γ response, we evaluated which markers are predictive of infection with pH1N1 vs. H3N2. Expression of CD112, PANHLA, CD54, HLA-C, and MICA predict pH1N1 infection, while CD111 (Nectin-1), CD95, and IP-10 predict H3N2 infection (**Fig. 4E**). PCA visualization indicates that expression of CD112 and CD54 are the major contributors to variance between pH1N1 and H3N2 infection (**Fig. 4F**). Ligand expression on infected pure monocytes at 24 HPI hours is displayed in **Fig. S10**, and is similar to those observed in the monocyte – NK cell co-culture, with CD112 and CD54 contributing strongly to strain variance (**Fig. S10F**), indicating that the strain-specific modulation of these ligands occurs independently of NK cell cross-talk.

We performed several quality checks of the data and our analyses. We found that the number of cells evaluated is not a significant contributor to variance and does not distinguish populations of mock vs. infected cells at 24 HPI, either in NK cell co-culture or purified (**Fig. S11-12A, C, E**). Similarly, the individual donors do not drive the variance observed in mock vs. infected cells or pH1N1 vs. H3N2 infection (**Fig. S11-12, B, D, F**). Thus, these data indicate a pattern of influenza infection across multiple healthy donors leading to changes in inhibitory and activating ligand expression, with expression of the activating ligand CD112 and the cell adhesion molecule CD54 distinguishing pH1N1 from H3N2 infection.

### Contribution of CD112 and CD54 to strain-specific NK cell IFN-γ production

Consistent with our mass cytometry and GLMM results, fluorescence cytometry corroborates that both CD112 and CD54 are significantly downregulated by influenza infection, but that the frequency of CD112 and CD54-expressing cells is significantly higher on pH1N1-infected monocytes than on H3N2-infected monocytes (**Fig. 5A**). Both strains similarly downregulate the NK cell activating ligand CD155 and while upregulating CD48 (**Fig. 5A**). To identify whether infection directly contributes to the modulation of CD112 and CD54 ligand expression, we examined whether the infected cells, identified as influenza nucleoprotein (Flu-NP)^+^, differentially expressed NK cell ligands. CD112 expression is predictive of the uninfected bystander (Flu-NP^-^) monocytes after exposure to H3N2 or pH1N1 infection (**Fig. 5B-C; underlined),** while CD54 expression is predictive of infected monocytes (Flu-NP^+^) following both H3N2 and pH1N1 infection (**Fig. 5B-C; underlined**). These results were confirmed by conventional flow cytometry (**Fig. 5D-E**). This pattern of ligand expression is independent of infection levels (**Fig. 5F-G**). Collectively, these data indicate that H3N2 infection leads to a more robust downregulation of CD112 and CD54 than pH1N1 infection; CD112 expression is retained preferentially on uninfected bystander (Flu-NP^-^) cells, while CD54 expression is retained on infected (Flu-NP^+^) cells (**Fig. 5D-G**).

**Fig. 5.**
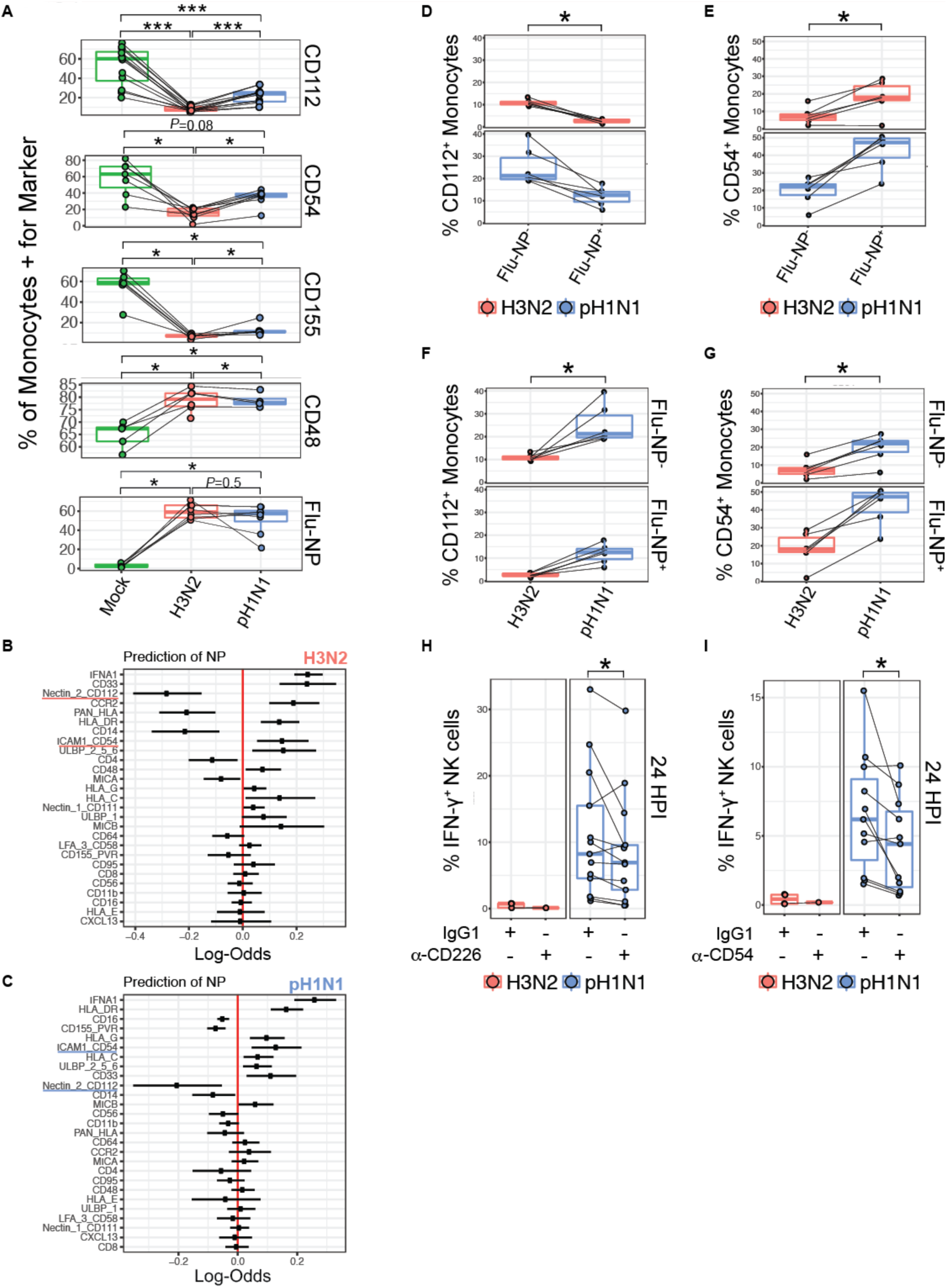
CD226 and CD54 contribute to strain-specific NK cell IFN-γ production in a subset of donors. **(A)** Expression of CD112, CD54, CD155, CD48 and Flu-NP by fluorescence flow cytometry at 24 HPI. GLMM prediction of H3N2- (**B**) or pH1N1- (**C**) exposed Flu-NP^-^ vs. Flu-NP^+^ monocytes at 24 HPI. **(D-G)** The frequency of CD112 and CD54 expression on infected (Flu-NP^+^) and uninfected (Flu-NP^-^) monocytes exposed to H3N2 or pH1N1 at an MOI of 3; *n*=6. **(H)** and **(I)** Frequency of IFN-γ^+^ NK cells in co-culture with influenza-infected monocytes following pre-incubation with saturating concentrations of isotype control or blocking antibodies to CD226, *n*=12 **(H)**, or CD54, *n*=11 **(I)**. \**P* < 0.05, \*\*\**P* < 0.0005, Wilcoxon signed-rank test.

Engagement of CD112 triggers NK cell activation through CD226 *(39)*. A blocking antibody to CD226 significantly reduces the NK cell IFN-γ response to pH1N1 compared to isotype control treatment (*P*=0.05), while the IFN-γ response to H3N2 remains near baseline at 24 HPI (**Fig. 5H**). This suggests that CD226 plays a role in the strain-specific IFN-γ response to pH1N1, though there is variation between individuals in their reliance on CD226. Similarly, inhibiting the interaction between CD54 and its cognate receptor LFA-1 with a CD54 blocking antibody *(40, 41)* significantly dampens the NK cell IFN-γ response to pH1N1 (*P*=0.020) (**Fig. 5I**). Though CD48, the cognate ligand for 2B4, is significantly elevated by H3N2 and pH1N1 infection, blockade of the CD48:2B4 interaction *(42)* does not consistently alter NK cell IFN-γ secretion in response to pH1N1 (**Fig. S13A**). The NKG2D ligand MICA is modestly predictive of pH1N1 infection vs. H3N2 at 24 HPI (**Fig. 4C; underlined**). Yet despite a prior report that NKG2D is involved in influenza recognition of A/Panama/2007/1999 (*35*), NKG2D ligands are not significantly upregulated by pH1N1 or H3N2, nor does NKG2D blockade significantly alter NK cell IFN-γ secretion in response to pH1N1 infection (**Fig. S13**). Collectively, these data indicate that CD226:CD112 and LFA-1:CD54 interactions play a critical role in triggering the sustained NK cell IFN-γ response to pH1N1, which is not observed with H3N2.

### Transcriptional profiling of pH1N1 and H3N2-infected monocytes

The incomplete inhibition of the NK cell IFN-γ response after blocking receptor ligand interactions suggests that additional factors are required. Thus, we performed RNA sequencing (RNA-seq) on mock-treated or infected monocytes in which we matched infection levels between pH1N1 and H3N2 (**Fig. 6A**). PCA reveals that the effects of infection and time point are far greater than the effects of individual donors, and that the two viruses have distinguishable transcriptional signatures (**Fig. 6B-D**). The genes differentially expressed between pH1N1 and H3N2 infection were imported into STRING, a database of protein-protein interactions paired with interaction confidence scores to construct an interaction network that stratifies the subtype-specific response (**Fig. 6E**) *(43, 44)*. When placed into this context at a false discovery rate of 0.01, a large number of transcripts that are significantly elevated by pH1N1 compared to H3N2 infection form a module of interactions connected by an interferon-α/β receptor (*IFNAR1*) node (**Fig. 6E and Table S2**). Numerous interferon-α related genes are upregulated in pH1N1-infected monocytes compared to H3N2, including *IFNA1* at 7 HPI, indicative of an early strain-specific transcriptional response (**Fig. 6E; *IFNA1* inset**). Thus, while *IFNAR1* was itself not significantly differentially expressed, this framework identified this module as a potential key difference in the enhanced and sustained NK cell IFN-γ response to pH1N1 compared to H3N2 infection. RNA transcript levels of *NECTIN2* (CD112) and *ICAM1* (CD54) are higher in pH1N1-infected cells compared to H3N2 infected cells, levels of *PVR* (CD155) RNA are similarly downregulated by both strains, and *CD48* RNA levels are elevated by infection, all in agreement with our observations at the protein level (**Fig. S14A-D**). Influenza RNA transcripts levels are similar between strains, though levels of the nuclear export protein are lower in H3N2 infection (**Fig. S14E**). To capture additional strain-specific protein-protein interactions, a network subgraph with lower confidence intervals (FDR = 0.1) was generated (**Fig. S14F**); gene names and corresponding p-values for 0.01 and 0.1 false discovery rates are listed in **Table S2**.

**Fig. 6.**
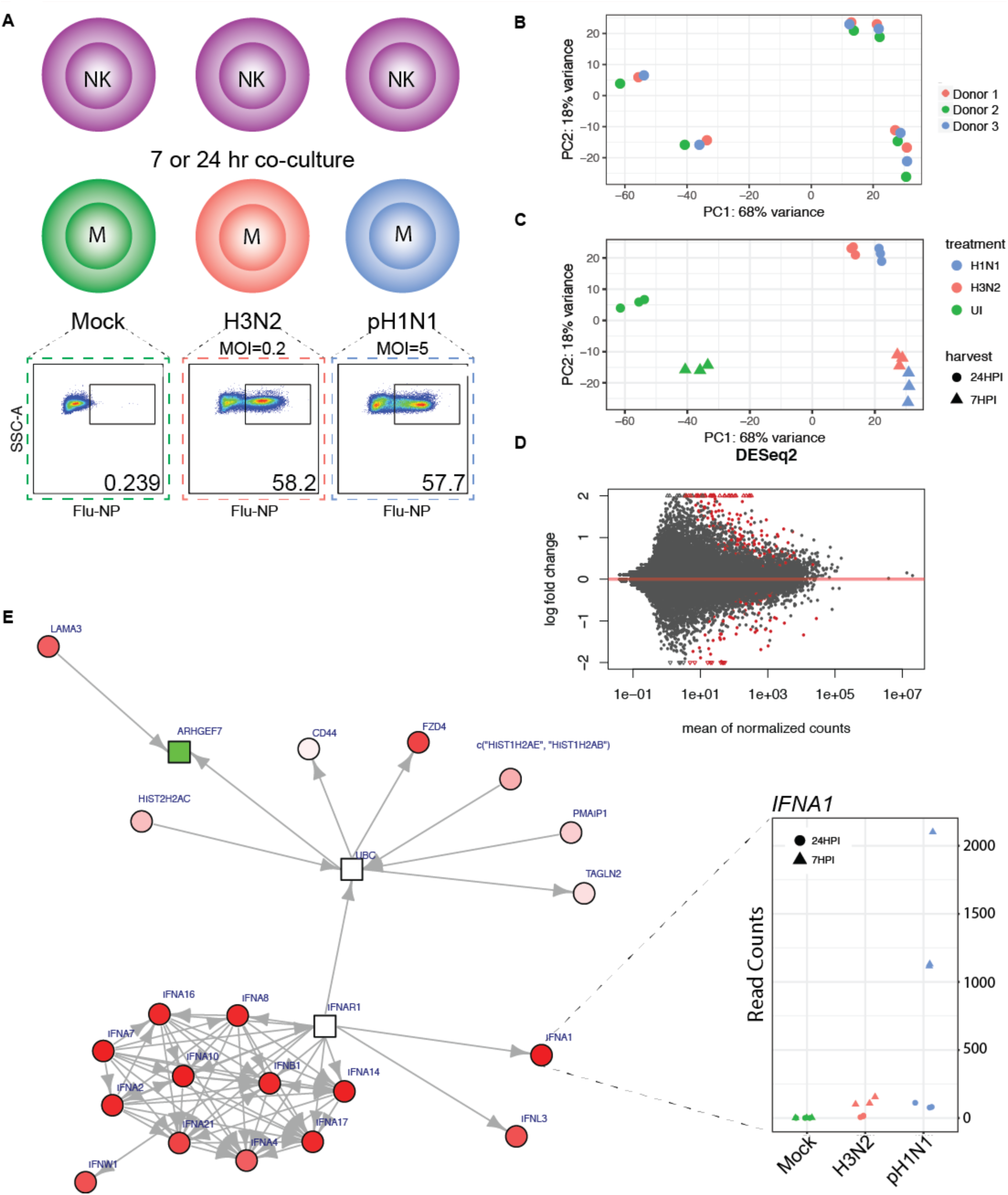
Whole transcriptome profiling of mock-treated, or pH1N1- or H3N2-infected monocytes reveals strain-specific alterations in the magnitude of the IFN-α response. **(A)** Schematic of RNA-sequencing experimental design. Monocytes from three healthy blood bank donors were mock treated or infected with pH1N1 (MOI=5) or H3N2 (MOI=0.2), which leads to approximately equal levels of Flu-NP^+^ monocytes followed by NK cell co-culture. 7 or 24 HPI, monocytes were re-isolated by magnetic separation followed by RNA extraction, cDNA library preparation and paired-end illumina sequencing. **(B)** PCA plot of the samples colored by treatment. (**C)** PCA plot of the samples colored by donor. **(D)** Volcano plot of differentially expressed genes between pH1N1 and H3N2 infected monocytes generated by DeSeq2, with the log2 fold change of each gene plotted against the total number of counts recorded for that gene. Differentially expressed genes with a p-value < 0.05 are highlighted in red. Triangles represent data points outside the graph area. **(E)** Differentially expressed genes were mapped to known protein-protein networks derived from the human STRING database. Individual gene are nodes in the network with scores assigned to them derived from a beta-uniform mixture model fitted to the unadjusted p-value distribution accounting for multiple testing. A subgraph (false discovery rate = 0.01) was generated with differential gene expression colored in red (up-regulated; pH1N1 > H3N2), green (down-regulated; pH1N1 < H3N2) or white (neutral). Shapes represent scores: rectangles are negative and circles are positive. Red cluster in bottom left: type 1 interferon-mediated signaling pathway elevated in pH1N1-vs. H3N2-infected monocytes. Inset: *IFNA1* mRNA counts within each condition.

### Strain-specific NK cell reactivity is modulated by exposure to differential cytokine production by infected target cells

In light of the strain-specific differences in transcripts associated with the cytokine IFN-α, we compared cytokine concentrations in supernatants from pH1N1 and H3N2 infected cells. pH1N1-infected monocytes secrete greater levels of many cytokines and chemokines, including IFN-α and several others associated with enhanced NK cell anti-viral responses including IL-12, IL-15, and IL-18 (**Fig. 7A and Table S3**). Blocking the activity of IFN-α using a monoclonal antibody specific to IFN-α receptor chain 2 (IFNAR2) significantly reduces IFN-γ secretion by NK cells, as does blockade of IL-15, IL18, and IFNGR1, but not IL-12 (**Fig. 7B**). Thus, signaling through IFN-α, IL-15, IL-18 and IFN-γ receptors are critical to the NK cell IFN-γ response to pH1N1. To test whether receptor-ligand interactions function cooperatively with the interferon-α receptor for the NK cell response to influenza, we assessed whether co-blockade of CD226 or CD54 and the IFN-α receptor leads to dampening of the NK cell IFN-γ response. Indeed, co-blockade of the CD226 with IFN-α receptor or CD54 with IFN-α receptor leads to further impairment in NK cell IFN-γ production, reducing IFN-γ to background levels (**Fig. 7C**), and accounting for almost the entirety of the strain-specific response.

**Fig. 7.**
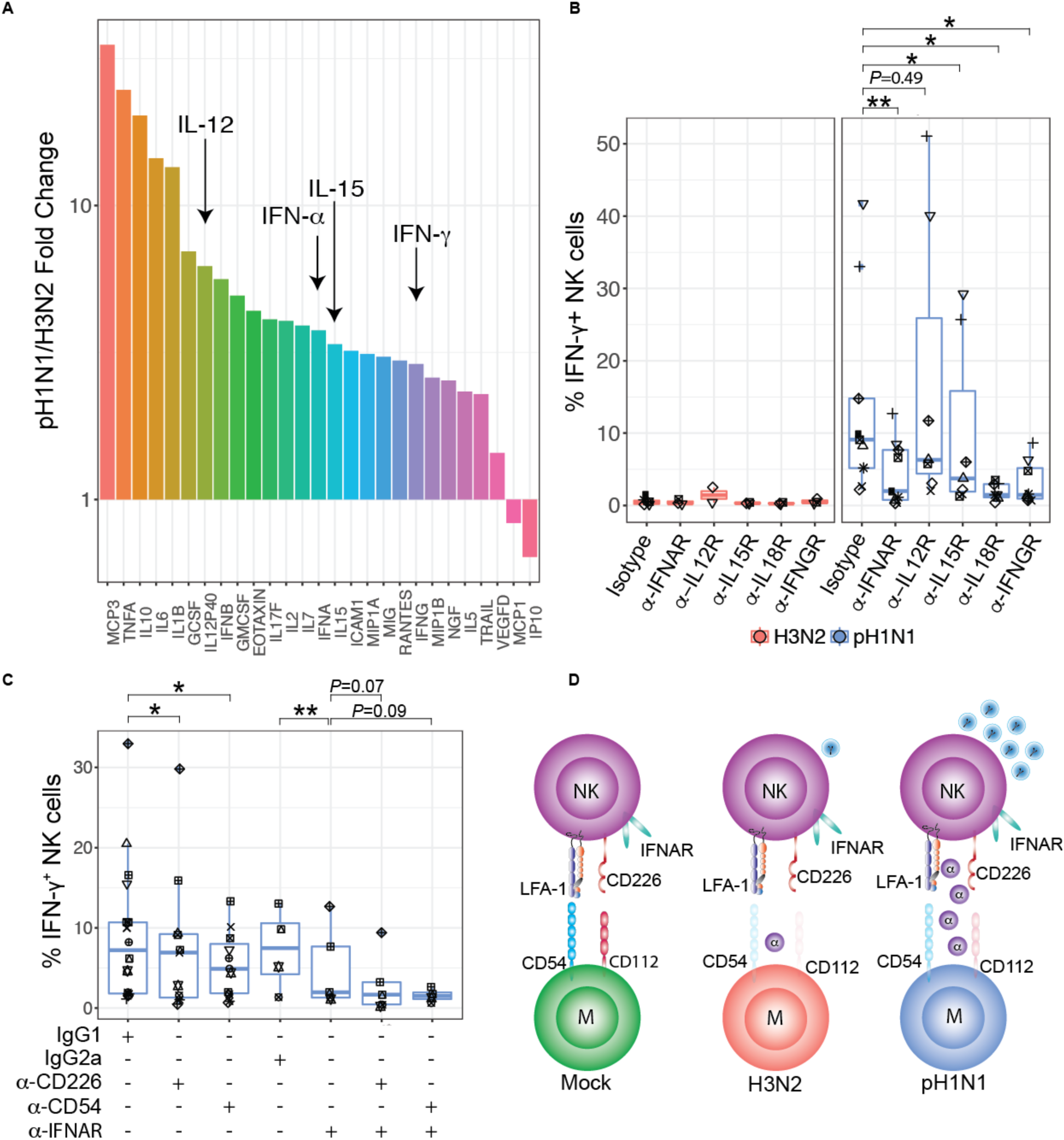
Cytokine profiles of pH1N1- and H3N2-infected monocytes contribute to strain-specific NK cell IFN-γ response. **(A)** Cytokine concentrations assessed by Luminex®, values displayed represent the mean fluorescence intensity in pH1N1-infected/H3N2 infected conditions (MOI=3). Cytokines elevated by 2.5-fold over the level in mock-infected monocytes are plotted. **(B)** Impact of cytokine receptor blocking on the NK cell IFN-γ response evaluated by pre-incubating NK cells for 1 hr with blocking antibodies specific to IFNAR2, IL-12R, IL-15R, IL-18R, and IFNGR1 followed by co-culture with infected monocytes. 24 HPI, intracellular cytokine staining was used to assess NK cell IFN-γ^+^ frequency compared to treatment with an isotype control antibody (H3N2: *n*=2-6, pH1N1: *n*=9). **(C)** NK cells were incubated for 1 hr with antibodies specific to IFNAR2 and CD226 or CD54 followed by co-culture with autologous infected monocytes. At 24 HPI, intracellular cytokine staining was used to assess IFN-γ production compared to treatment with an isotype control antibody (*n*=6). \**P* < 0.05, \*\**P* < 0.005, Wilcoxon signed-rank test. **(D)** Model of strain-specific NK cell recognition of influenza A infection. pH1N1-infection of monocytes virus does not downregulate CD54 and CD112 to the same extent as H3N2-infected monocytes. CD54 expression is retained on Flu-NP^+^ cells while CD112 expression is preferentially retained on exposed, uninfected monocytes. pH1N1 infection of monocytes elicits enhanced IFN-α secretion and blockade of IFNAR2 dampens NK cell anti-pH1N1 IFN-γ production.

## Discussion

Our study defines a previously unrecognized pathway whereby NK cells, despite lacking antigen-specific receptors such as those expressed by B and T cells, exhibit pathogen specificity. We report that antigenically divergent influenza strains trigger NK cells to produce a distinct functional response based upon the strain-specific downregulation of CD112 and CD54 and changes in IFN-α production (**Fig. 7D**). The pH1N1 strain stimulates higher IFN-α production and fails to fully downregulate activating ligands for NK cells, leading to a dramatically greater and more sustained IFN-γ response compared to the H3N2 strain. This pathway, which involves differences in the induction of cytokines production and ligands in response to distinct viruses, provides the first insight into the cellular mechanisms responsible for tuned innate responses to specific viral strains in human NK cells. Such differences could significantly affect disease pathogenesis during *in vivo* infection, when the need to clear the virus is balanced by the threat posed by an over-exuberant immune response, which is strongly implicated in influenza mortality *(45, 46)*.

The abrogation of IFN-γ production upon loss of direct contact between NK cells and infected target cells implicates receptor-ligand involvement in the NK cell recognition of infected cells. Further, based on earlier reports that influenza A infection leads to the upregulation and re-organization of HLA class I molecules *(47)*, we hypothesized that activating receptors may play a more important role in the recognition of influenza-infected cells than ‘missing self’. Indeed, our data indicate that HLA class I expression is retained on influenza-infected cells, highlighting how retention or even upregulation of HLA class I molecules on influenza-infected cells is insufficient to shield from NK cell recognition. Therefore, we were initially surprised to discover that the NK cell activating ligands CD112 and CD54 are expressed at *lower* levels on infected than uninfected cells. Despite their lower levels, these ligands clearly play a critical role in recognition of influenza-infected cells, as blockade of CD112:CD226 or CD54:LFA-1 interactions significantly diminishes IFN-γ production in response to the pH1N1 strain.

Our data suggest that influenza downregulates ligands for activating receptors as an NK cell immunoevasion strategy. Intriguingly, the strain specificity in the magnitude of activating ligand downregulation suggests that H3N2 strain has evolved to escape NK cell recognition by thoroughly downregulating expression of CD112 and CD54, while the pH1N1 strain does so incompletely, leading to more robust detection and response. Infection of mouse alveolar epithelial cells with influenza virus (A/PR/8/34 (H1N1)) induces expression of CD54 *(48)*, indicating the influenza-mediated modulation of CD54 occurs in an *in vivo* model of infection. Finally, though CD112 and CD54 emerged as the dominant factors contributing to the strain-specific variance in IFN-γ production in our CyTOF data set, additional receptors are almost certainly involved. For instance, the 2B4 ligand CD48 is significantly predictive of H3N2 infection, and may contribute to recognition, at least in a subset of individuals. Further, MICA significantly predicts infection of pH1N1, suggesting that its cognate receptor NKG2D may play a role in recognition, even if its role can be compensated by additional ligands, as blockade does not significantly diminish NK cell IFN-γ responses pH1N1 (**Fig. S13B-C**).

While a role for the CD226:CD112 interaction in NK cell recognition of influenza-infected cells has not been reported previously, other viruses modulate CD112. For instance, the gD glycoprotein of alpha-herpes simplex virus-2 (HSV-2) downregulates CD112, reducing CD226 binding and NK cell-mediated lysis of HSV-2-infected cells *(49)*. Further, CD112 and CD155 are downregulated by cytomegalovirus (CMV) in dendritic cells, and CD226 blockade impairs NK cell responses to CMV *(50, 51)*. In HIV-1, infected primary CD4^+^T cells express CD155 and trigger NK cell-mediated lysis of the infected cells by through CD226 *(52)*. In our data, CD155 (PVR) expression is predictive of mock infection and downregulated robustly by H3N2 and pH1N1 (**Fig. 4-5**). Collectively, viral modulation of CD226 ligands is a viral strategy employed recurrently to evade NK cell recognition, highlighting the evolutionary pressure placed by human NK cells on influenza A viruses. It is intriguing that pH1N1, the strain most recently introduced into the human population, displays the least ability to escape from NK cell recognition.

While the NK cell IFN-γ response is clearly dependent on receptor-ligand interactions, we find that production of inflammatory cytokines by infected cells is also necessary, but not sufficient, to trigger full NK cell activation. Blockade of IFNAR2, IL-15R, IL-18R and IFNGR1 robustly restricts NK cell IFN-γ production when cell physical contact is preserved. These data are consistent with NK cells requiring multiple signals provided by IFN-α (or certain other cytokines), CD112, and CD54 for IFN-γ secretion. Prior work found that NK cell responses to dengue virus involve type I IFN, TNF and HLA class I downregulation, providing a distinctive example in which the two-signal requirement is met by activating ligands and loss of inhibitory signals (*8*). Beyond viral infection, Bryceson and colleagues have shown that while cross-linking CD16 induces NK cell activation, signaling through any other one receptor is insufficient to activate NK cells, requiring instead engagement of specific pairs of NKp46, NKG2D, CD226, CD244 or CD2 *(53, 54)*. Collectively, our data are consistent with a “two-signal” model, in which a specific cytokine milieu acts to sensitize NK cells, which in turns confers susceptibility to activation through even low levels of activating ligands that are retained on the surface of infected cells.

Here we found that two highly expressed NK cell-activating receptors govern the quality and durability of the NK cell response. Human primary NK cells are consistently >70%^+^ for CD226 or LFA-1 expression *(42)*, yet the magnitude of the NK cell response to pH1N1 or H3N2 differs considerably between individuals, calling into question the source of this unique activation profile among individuals. One possibility is that monocytes from individual donors produce highly variable levels of inflammatory cytokines post-infection, thus eliciting differential NK cell priming. Another possibility is that donor vaccination and infection history may influence intrinsic NK cell activity to *in vitro* influenza infection. Specifically, in light of recent evidence supporting NK cell memory responses in mice, primates, and humans *(55-60)*, it is possible that prior exposure could have shaped memory-like responses in our cohort, leading to differential responses based on immunization and infection history. Indeed, using a human cohort study of individuals vaccinated with pH1N1, NK cell IFN-γ responses were elevated four weeks post-vaccination and this enhancement was dependent on IL-2 and influenced by human CMV seroconversion status *(31)*. Intriguingly, NK cell reactivity measured four weeks post-vaccination was also elevated in response to innate cytokines and this capacity persisted up to six months *(31, 32)*. Along similar lines, the molecular basis for stable IFN-γ expression in human cytomegalovirus-specific NK cell populations was traced to epigenetic remodeling of the noncoding region upstream of the *IFNG* locus *(61)*. It is unclear if similar epigenomic changes govern memory-like responses to influenza. Investigation of influenza-naïve human NK cells would be informative; however, it is problematic to obtain reliably influenza naïve individuals. For instance, in one study, 100% of the children evaluated had antibodies to at least one influenza strain by age six *(62)*. Further, though cord blood may be a source of naïve NK cells, they are not fully functional *(63, 64)*.

The viral characteristics responsible for strain-specific modulation of cellular NK cell receptor ligands and cytokine production remain unknown. A higher replicative fitness of H3N2 vs. pH1N1 in monocytes could imbue H3N2-infected cells with additional copies of NS1, the main type I IFN antagonist encoded by influenza A viruses *(65-67)*, accounting for part of the strain differences. We suspect NS1 protein copy number may not be the sole contributing factor, as NK cell IFN-γ production did not inversely correlate with infection levels when NK cell activity was tested against four additional influenza strains and NS1 transcript levels were also similar (**Fig. 1 and S14**). Further, when a higher titer of pH1N1 was used to achieve similar infection levels, the NK cell IFN-γ production remained significantly higher for pH1N1 compared to H3N2 (**Fig. S4**). Another possibility is variation in the NS1 sequence (82% sequence identity) may enable H3N2 to disarm the host type I IFN system with higher efficiency than pH1N1. Strain-specific NS1 activity is observed between the NS1 from the 1918 pandemic influenza strain vs. A/WSN/33, with pandemic NS1 more efficiently blocking expression of IFN-regulated genes *(68)*. Whether sequence divergence and/or protein copy number in NS1 or other genes between influenza A strains contributes to differential NK cell activation is a key future endeavor.

This study applied tailored statistical approaches to CyTOF and RNA-seq datasets to extract candidate proteins and genes to test for their role in triggering strain-specific NK cell responses. In our CyTOF analysis approach, the donor-specific variance was modeled using a GLMM to estimate parameters at the population level and remove the donor-specific variability. For CyTOF classification analyses, marker expression levels are explanatory variables and no explicit model assumptions about their distribution need be made. In the case of zero-inflated expressions, the estimated marker coefficients are shrunk towards zero and provide more conservative estimates. Furthermore, multimodal distributional markers do not violate the logistic likelihood model assumptions. In our regression analyses, we assume a Gaussian likelihood model for the response marker, i.e. Flu-NP. However, if these markers are highly multi-modal, then the Gaussian likelihood model would not be appropriate. To investigate this issue, we evaluated Poisson and quasi-Poisson likelihood models. We observe consistent estimates across all three models and conclude that our results depend only weakly on the choice of the model. Further, we used PCA to visualize our CyTOF results and overlaid original variables onto the plots to interpret principal components in terms of markers. While PCA may provide suboptimal results for data that is multimodal or zero-inflated, the PCA plots are consistent with our GLMM findings, corroborating CD54 and CD112 as major drivers of differential expression between H3N2 and pH1N1 influenza strains.

For RNA-seq analyses, prior knowledge of protein-to-protein interactions networks from the STRING database was incorporated by placing p-values onto known networks, allowing the construction of transcript subgraphs displaying differentially expressed genes *(43)*. Such analysis allows for the inclusion of genes not significant by themselves but provide an important linking structure connecting differentially expressed gene clusters. In our analysis, we observed that *IFNAR1* links a network of interferons and *IFNA1* is differentially expressed in pH1N1 vs. H3N2.

A limitation of our study is the use of human monocytes in lieu of lung epithelial cells, which are the primary influenza target cells during natural infection. The use of blood monocytes allows for the maintenance KIR and HLA class I ligand interactions, preventing NK cell responses due to KIR-HLA class I mismatch rather than infection. This is an important consideration as KIR and HLA class I have been found to play a critical role in the outcome of several viral infections *(69-71)*. For instance, the KIR2DL3^+^/HLA-C*03:04 association with the T_gag303_V mutation of HIV-1 leads to enhanced binding and inhibition of KIR2DL3^+^ NK cell degranulation *(70-72)*.

This work highlights how differences in innate responsiveness to distinct viral strains may significantly impact the balance between protection and immunopathology *in vivo*. Consistent with this idea, pH1N1 leads to higher induction of pro-inflammatory cytokines in the lungs of ferrets and macaques compared to infection with seasonal A/Kawasaki/UTK-4/09 (H1N1) at three days post-infection *(73)*. Collectively, these data raise the intriguing possibility f calibrating NK cell response through modulation of the cytokine environment and expression of activating ligands. Along these lines, the proteasome inhibitor bortezomib enhances NK cell killing of multiple myeloma cells by upregulating NKG2D and CD226 ligands on tumor cells *in vitro (74)*. Whether such approaches could be similarly leveraged to shape the quality of the innate and subsequent adaptive response to infection or vaccination *in vivo* will be an important future endeavor.

## Materials and Methods

### Study Design

For all experiments, leukoreduction system chambers were purchased from the Stanford Blood bank. As subjects were fully de-identified, the study protocol was deemed not to be human subjects research by the Institutional Review Board at Stanford University. All experiments were performed on PBMCs separated using Ficoll-Pacque (GE Healthcare) density gradient centrifugation from whole blood and cryopreserved in 90% (vol/vol) FBS (Thermo Scientific) plus 10% (vol/vol) dimethyl sulfoxide (Sigma-Aldrich).

### Virus Production and Titration

A/Victoria/361/2011 (H3N2), A/California/07/2009 (pH1N1), A/Bayern/7/1995 (H1N1) (Influenza Reagent Resource), A/Beijing/32/1992), A/Hong Kong/1/68 (H3N2), and A/Melbourne/1/1946 (H1N1) were grown in 10-day-old embryonated specific pathogen-free chicken eggs (Charles River Laboratories International, Inc.) at 37°C and 55-65% humidity. Allantoic fluid was harvested 48 hours post-infection. A/Puerto Rico/8/34 (H1N1) was grown in Madin-Darby canine kidney (MDCK) cells. Viral titer was determined by plaque assay on MDCK cells in the presence of 2 μg/mL L-1-tosylamido-2-phenylethyl chloromethyl ketone (TPCK)-treated trypsin.

### Cell purification, Stimulation, and Infection

Cryopreserved PBMCs were thawed and washed with RPMI (Thermo Scientific) supplemented with 10% (vol/vol) FBS (Thermo Scientific) and 50 U/mL benzonase (EMD Millipore). NK cells and monocytes were purified by magnetic-activated cell sorting via negative selection (Miltenyi). PBMCs were cultured in RPMI 1640 (Invitrogen) supplemented with 10% fetal bovine serum (FBS) + 2 mM L-glutamine + antibiotics [penicillin (100 U/ml), streptomycin (100 mg/ml); Life Technologies] at 37°C with 5% CO_2_. For influenza infections, monocytes were washed and re-suspended in serum-free media at 0.75 × 10^6^ per 100 μL and infected at a multiplicity of infection (MOI) of 3 for 1 h at 37°C with 5% carbon dioxide. 1 HPI, viral inoculum was removed and cells were re-suspended in 200 μL of RPMI supplemented with 10% FBS + antibiotics [penicillin (100 U/ml), streptomycin (100 mg/ml) with autologous NK cells. After a further 2 or 19 hour incubation, 2 μM monensin, 3 μg/mL brefeldin A (eBiosciences), and anti-CD107a-allophycocyanin-H7 (BD Pharmingen) were added to the co-culture for 4 hours. EDTA (Hoefer) was added at a final concentration of 1.66 mM for 10 min to arrest stimulation. In *trans*-well experiments, NK cells and monocytes were separated during culture by a semipermeable membrane of 0.4-μm size pore (Corning).

### Flow Cytometry

Cells were stained with LIVE/DEAD fixable Aqua Stain (Life Technologies), followed by surface staining with anti-CD56-phycoerythrin-Cy7, and then fixed and permeabilized with FACS Lyse and FACS Perm II (BD Pharmingen) according to the manufacturer’s instructions. Cells were stained with anti-CD3-phycoerythrin (clone: OKT3; BioLegend), anti-CD16-PerCPCy5.5 (clone: 3G8; BioLegend), anti-IFN-γ -violet-450 (clone: B27; BD Biosciences), antinucleoprotein-fluorescein isothiocyanate (clone: AA5H; abcam)) and anti-CD7-allophycocyanin (clone: 6B7; BioLegend), or anti-CD14-allophycocyanin (clone: HCD14; BioLegend) and fixed using 1% paraformaldehyde. Ligand staining was performed after blocking in Human TruStain FcX^TM^ Fc receptor blocking solution (BioLegend) according to the manufactures instructions followed by staining with CD54-PE (clone: HA58; BioLegend), CD155-PE (clone: SKII.4; BioLegend), CD112-PE (clone: Tx31; BioLegend) or CD48-PE (clone: BJ48; BioLegend), ULBP2/5/6 (clone: 165903; R&D Systems), ULBP1 (clone: 170818; R&D Systems), MICA (clone: 159227; R&D Systems), MICB (clone: 236511; R&D Systems). Uncompensated data were collected using a three-laser MACSQuant® Analyser (Miltenyi Biotec). Analysis and compensation were performed using FlowJo^TM^ flow-cytometric analysis software, version 9.9.4 (Tree Star). Fluorescently activated cell sorting was performed using a BD FACS Aria instrument. Viral suppression was calculated as [% Flu-NP^+^ in pure monocytes - % Flu-NP^+^ in monocytes + NK cells) / (% Flu-NP^+^ in pure monocytes) x 100].

### RNA isolation and RT-qPCR

RNA was isolated at the indicated time points from either co-culture (RT-qPCR) or re-isolated NK cells or monocytes (RNA-seq) using RNA-Bee (Tel-Test) according to the manufacturer’s instructions. For RT-qPCR, total RNA was reverse transcribed using SuperScript® VILO^TM^ cDNA synthesis kit (Thermo Scientific) followed by triplicate qPCR reactions using *IFNG* (*IFNG* Taqman Gene Expression Assay, Life Technologies 4331182) or Flu-Matrix transcript-specific primers to an identical region in the pH1N1 and H3N2 strains with the TaqMan Universal Master Mix II on an StepOne^TM^ Plus Real-Time PCR system (Applied Biosytems) (Matrix primers Forward: 5’ - CTT CTA ACC GAG GTC GAA ACG TA-3’, Reverse: 5’-TTG GTC TTG TCT TTA GCC ATT CCA-3’). Averaged triplicate levels of RTqPCR results were normalized to 18S or GAPDH average duplicate levels. Relative fold inductions were calculated using the ΔΔC_t_ formula.

### RNA sequencing and Statistical Analysis

RNA was extracted from monocytes re-isolated from co-culture using negative magnetic selection. RNA was quality checked by Agilent 2101 Bioanalyzer followed by TruSeq cDNA library preparation and paired-end 75-bp mRNA-seq RNA-sequencing on the Illumina NextSeq platform. The raw influenza sequencing reads were aligned using STAR *(75)* to the influenza genome of each strain downloaded from the Influenza Research Database (pH1N1, CY121687; H3N2, KJ942683). The raw human sequencing reads were aligned to the Genome Reference Consortium Human Build 38 genome, downloaded from the Illumina webserver. Alignment was performed using Bowtie2 *(76)* and TopHat *(77)* to produce a parsimonious transcriptome assembly. The transcriptome assembly was analyzed using DEseq2 (R package on Bioconductor) *(78)* and differentially expressed genes were mapped to known protein-protein networks derived from the STRING database *(43)* using BioNet (R package on Bioconductor) *(44, 79)*. A low confidence thresholding increases the network size, lowering the risk of missing important subgraphs in the downstream analysis at the cost of an increased number of false positives. Individual genes are nodes in the network with assigned scores derived from a beta-uniform mixture model fitted to the unadjusted p-value distribution to account for multiple testing.

### Cytokine Multiplex Assay

The concentration of cytokines in supernatants following cell stimulation was assessed in duplicate using the cytokine multiplex technology by Luminex® according to manufacturer’s instructions using 100 μl of neat supernatant.

### Staining and Mass Cytometry Acquisition

Antibodies were conjugated using Maxpar X8 labeling kits (DVS Sciences) and lyophilized (Biolyph, LLC), with the exception of CD19-Qdot® 655 conjugate. Detailed staining protocols have been described *(80-83)*. Briefly, cells were live-dead stained with 25 μM cisplatin (Enzo Life Sciences), followed by surface staining for 30 min at 4°C with rehydrated lyospheres, fixed, permeabilized, and stained with rehydrated lyospheres containing the intracellular antibodies for 45 min at 4°C *(83, 84)*. The staining panel is listed in **Table S1**. Cells were fixed overnight in iridium-191/193 interchelator (DVS Sciences) and washed 1× in PBS and 3× in Milli-Q® H_2_O before acquisition on a Helios mass cytometer (Fluidigm Corp., South San Francisco). The data was normalized using EQ^TM^ Four Element Calibration Beads (Fluidigm Corp., South San Francisco) *(85)*. Lineage gating was performed using FlowJo^TM^ flowcytometric analysis software, version 10.2 (Tree Star).

### Receptor Blocking Tests

NK cells were pre-incubated with mAbs antibodies specific to CD226 (Clone 11A8, BioLegend), CD54 (Clone: BBIG-I1 (11C81), R&D Systems), purified mouse anti-human NKG2D (Clone 1D11, BD Pharmingen), 2B4/CD244 (Clone: PP35, eBioscience) at a final concentration of 10 μg/mL or with purified mouse IgG1, κ isotype control (BioLegend, IgG1 clone: MOPC-21, IgG2a clone: MG2a-53, 10 ug/mL) for 1 h prior to co-culture. The IFNAR chain 2 (PBL Biomedical Laboratories, clone: MMHAR-2) and Human IL-18 R alpha/IL-1 R5 Antibody (R&D Systems, MAB840-SP) were used at a final concentration at 2.5 μg/ml. IL-12R *β* 2 Ab (R&D Systems AF1959-SP) was used at a final concentration of 83.33 μg/mL. Human IL-15R alpha antibody was used at a final concentration of 4 μg/mL (Santa Cruz, sc-374023). Mouse anti-human CD119 IFN-γ receptor α chain (Clone: GIR-208, BD Pharmingen) was used at a final concentration of 50 μg/mL.

### Statistical analysis of CyTOF data

Statistical testing was performed in the open-source statistical programming language R *(86)*. For mass cytometry analysis, the R packages glm and mbest were used. For our classification analyses, a logistic regression models was constructed to identify markers that were significantly associated with response variables. Cell infection status (i.e. mock, pH1N1 or H3N2) constituted the response variables, while antibodies were considered explanatory variables. Raw counts were transformed using an arcsinh transformation with a standard cofactor of 5. Estimated coefficients are in units of log-odds, which are logarithm of ratios of two probabilities: the probability that a cell is B (for example, pH1N1) (encoded as 1) over the probability that a cell is A (for example, H3N2) (encoded as 0). The logistic regression model for the *i*th cell is a linear combination of weighted (denoted by *β*) marker intensities (denoted by *x*):
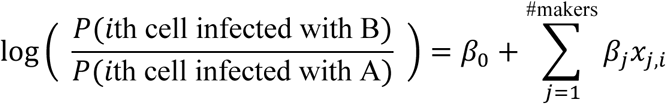

To account for donor specific variability, we used logistic regression with a random effects term (denoted by *z*):

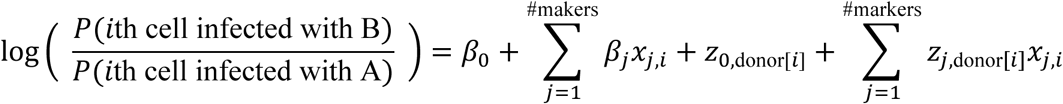

Markers with positive coefficients indicate that an increase in the associated marker increases the log-odds while markers with negative coefficients indicate that an increase in the associated marker decreases the log-odds. For our regression analyses, we used a Gaussian likelihood model with one marker as the response variable, e.g. Flu-NP, and all other markers as the explanatory variables.

## ACKNOWLEDGEMENTS

Thank you to Drs. Leslie Goo, Taia Wang, Cristina Tato and members of the Blish Lab for critical reading of the manuscript. We wish to thank Dr. Sara Prescott (Stanford University) for her assistance with the RNA-seq alignment to the human transcriptome and Rachel Hagey and Dr. Taia Wang for influenza strains. We acknowledge the Stanford Human Immune

Monitoring Center, in particular Dr. Yael Rosenberg-Hasson for the Luminex assay and Dr. Michael Leopold for assistance with the Helios mass cytometer. RNA sequencing was performed at the Stanford Functional Genomics Facility. Cell sorting was conducted on instruments in the Stanford Shared FACS Facility.

## Author contributions

Conceptualization: L.M.K and C.A.B.; Experiments: L.M.K and R.V. Investigation: L.M.K and C.S.; Writing: L.M.K., C.S., S.P.H., and C.A.B.; Funding C.A.B and S.P.H.

## Data and materials availability

All .fcs files will be uploaded into ImmPort (Accession numbers #) and RNA-seq into GEO (Accession numbers #).

## Supplementary Materials

**Fig. S1.** Flow cytometry gating scheme and infection levels of autologous monocyte – NK cell co-culture system.

**Fig. S2.** NK cell suppress viral Flu-NP levels at 7 HPI and matrix RNA levels at 7, 24 and 48 HPI.

**Fig. S3.** NK cells produce IFN-γ in response to H1N1 infected monocytes after T cell depletion to <0.25%, and adding back T cells does not increase responses.

**Fig. S4.** Frequency IFN-γ^+^ NK cells after exposure to pH1N1 and H3N2 infected monocytes.

**Fig. S5.** Serial gating strategy to define monocytes by mass cytometry.

**Fig. S6.** Examples of CyTOF surface stains on monocytes stained with the NK cell ligand panel after mock, pH1N1, or H3N2 infection.

**Fig. S7.** Examples of CyTOF intracellular stains on monocytes after mock, pH1N1, or H3N2 infection.

**Fig. S8.** Identification of influenza-mediated modulation of NK cell inhibitory and activating ligands on monocytes co-cultured with NK cells for 6 hours using mass cytometry and GLMM analysis.

**Fig. S9.** Identification of influenza-mediated modulation of NK cell inhibitory and activating ligands on pure monocytes at 7 HPI using mass cytometry and GLMM analysis.

**Fig. S10.** Identification of influenza-mediated modulation of NK cell inhibitory and activating ligands on pure monocytes at 24 HPI using mass cytometry and GLMM analysis.

**Fig. S11.** Quality check of variables that could potentially influence the ability to identify makers predictive of infection in monocyte – NK cell co-culture.

**Fig. S12.** Quality check of variables that could potentially influence the ability to identify makers predictive of infection in pure monocyte culture.

**Fig. S13.** Role of 2B4 and NKG2D in NK cell recognition of mock, H3N2- and pH1N1-infected monocytes.

**Fig. S14.** Whole transcriptome profiling of mock-treated, or pH1N1- or H3N2-infected monocytes.

**Table S1.** Ligand CyTOF Antibody Panel.

**Table S2.** Genes differentially expressed between pH1N1 and H3N2 infected monocytes at a false discovery rate of 0.01 and 0.1.

**Table S3.** Mean Fluorescence Intensity Values of Raw Luminex Data.

